# LPS O-antigen polysaccharide length impacts outer membrane permeability of enteric gram-negative bacteria

**DOI:** 10.1101/2025.08.14.670410

**Authors:** Kerrie L. May, Tatsuya Akiyama, Bella G. Parker, Minsu Kim, Marcin Grabowicz

**Affiliations:** Department of Microbiology & Immunology, Emory University School of Medicine, Atlanta, GA 30322; Emory Antibiotic Resistance Center, Emory University School of Medicine, Atlanta, GA 30322; Department of Physics, Emory University, Atlanta, GA 30322; Division of Infectious Diseases, Department of Medicine, Emory University School of Medicine, Atlanta, GA 30322

## Abstract

The Gram-negative outer membrane (OM) forms the bacterial cell surface and acts as a barrier against antibiotic influx. In enteric species, the OM is covered by lipopolysaccharides (LPS) decorated with varying lengths of O-antigen (O-Ag) polysaccharide that protect bacteria against mammalian host defenses. Studies of lab-adapted *Escherichia coli* K-12 strains have proven instrumental in unravelling the essential processes of LPS synthesis, transport, and assembly into the OM. However, O-Ag synthesis was inactivated in K-12 strains during their lab adaption, and these cells produce a non-native, truncated LPS form. Surprisingly, we found that re-activating O-Ag synthesis in K-12 permeabilizes the OM to diverse antibiotics, causing susceptibility. The O-Ag that modifies LPS is directly responsible for the compromised OM barrier. Lengthening the O-Ag polysaccharide worsens antibiotic sensitivity; while shortening it, or removing it entirely, improves antibiotic resistance in both *E. coli* and the human pathogen, *Shigella flexneri*. Our data show that OM antibiotic barrier integrity is maintaining by a balanced production of long and short LPS forms, and that this balance is dysfunctional in model *E. coli* K-12 strains. Our findings reveal that long O-Ag polysaccharides are a double-edged sword: while well-recognized as critical for protection against external host assaults, their transport and assembly onto the surface comes at the inherent price of compromising the OM barrier. Hence, LPS production balances between competing needs in host defense and OM integrity. Moreover, we identify an inherent advantage for species that produce O-Ag-lacking lipooligosaccharide (LOS), rather than LPS.

**IMPORTANCE:** The outer membrane (OM) of bacteria like *Escherichia coli* and *Shigella flexneri* forms a barrier that protects cells against antibiotics and immune effectors. The surface-exposed leaflet is filled by lipopolysaccharides (LPS) decorated with long “O-antigen” (O-Ag) polysaccharides. The benefit of covering the surface with O-Ag is well-appreciated - these long polysaccharides shield against host assaults. Our study reveals a hidden cost to these long O-Ag polysaccharides: transporting and assembling LPS modified with O-Ag compromises integrity of the OM antibiotic barrier, rendering bacteria vulnerable to antibiotics. Cells must balance O-Ag across two parameters - protection from the host and preserving OM integrity. Our findings also present an inherent benefit to not producing O-Ag, a common feature among diverse bacterial pathogens.

## INTRODUCTION

The outer membrane (OM) of Gram-negative bacteria is an essential organelle that forms the cell surface and acts as a highly effective permeability barrier against toxins such as antibiotics. The OM is an asymmetric lipid bilayer with an inner leaflet composed of phospholipids (PLs) and an outer leaflet comprised of the glycolipid LPS (1). The chemical structure of LPS is tripartite: (i) “lipid A” is the membrane anchor to which (ii) a core oligosaccharide component is attached and is distally modified with (iii) a variable length O-antigen (O-Ag) poly- or oligo-saccharide (2, 3). Most Enterobacterales, such as *Escherichia coli* and *Shigella flexneri*, produce LPS molecules decorated with long O-Ag polysaccharides that extend far into the extracellular milieu (4). LPS modified with O-Ag is commonly called “smooth LPS”, whereas those lacking O-Ag are referred to as “rough LPS”.

LPS and O-Ag are both synthesized at the cytosolic face of the inner membrane (IM) but via distinct pathways (**Fig. 1A**). Lipid A is synthesized by the Raetz Pathway, and the core oligosaccharide is assembled during this process (3). After synthesis, LPS is flipped across the bilayer to the periplasmic leaflet (2, 3). O-Ag is independently synthesized on an undecaprenyl phosphate (Und-P) lipid carrier by a series of glycosyltransferases (the first committed step in *E. coli* is accomplished by the WbbL enzyme) (5, 6). Sugars are added sequentially to create a distinctly ordered oligosaccharide that forms the repeat unit of the O-Ag polysaccharide (5, 6). To form the polysaccharide, lipid-linked oligosaccharide units are flipped to the periplasmic leaflet of the IM and polymerized by the IM protein Wzy (5, 6). The length of O-Ag polysaccharides is characteristically modal (distributed within a particular size range) (5, 6). In many bacteria, including *E. coli* and *S. flexneri*, O-Ag modal length is defined by the IM protein Wzz (5, 6). The O-Ag polysaccharide is attached to the LPS core by the O-Ag “ligase” WaaL (**Fig. 1A**) (5, 6). The resulting smooth LPS molecules are crucial for virulence of pathogenic Enterobacterales, with rough LPS mutants being extensively characterized as severely attenuated in infection and pathogenesis models (7–17).

**Fig. 1.**
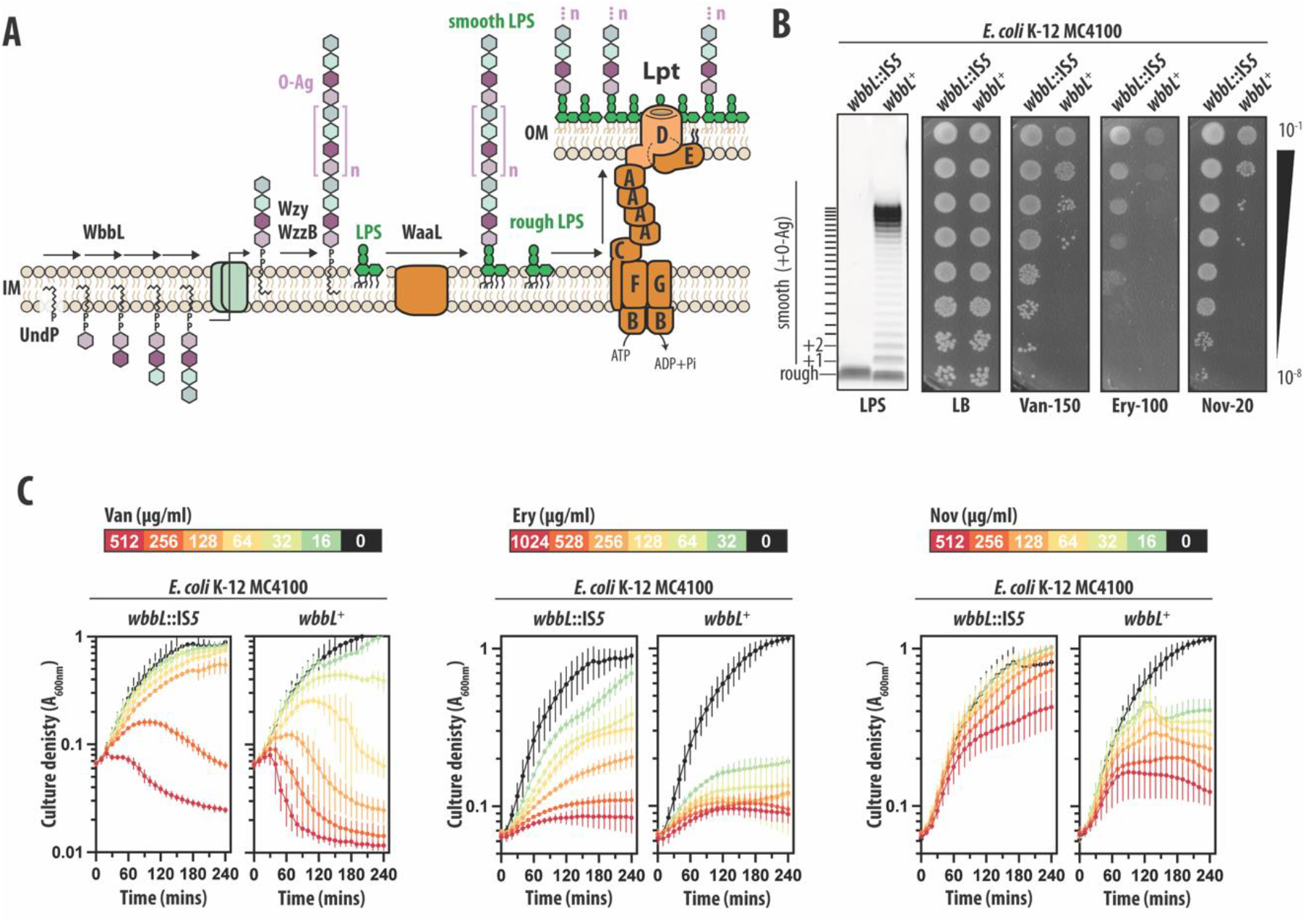
O-Ag production increases susceptibility of *E. coli* K-12 MC4100 to large-scaffold antibiotics. **(A)** Pathways for O-Ag biosynthesis, smooth LPS production, and LPS transport. **(B)** LPS profiles and antibiotic sensitivities of wildtype MC4100 (*wbbL*::IS*5*) and MC4100 *wbbL*^*+*^. Efficiency-of-plating assays with 10-fold serial dilution of cultures. **(C)** Time-kill curve analysis of vancomycin, erythromycin, and novobiocin sensitivity. All antibiotic concentrations are in μg/ml. All data are representative of at least three independent experiments.

Several decades of work, often in the genetically tractable *E. coli* K-12 strain MC4100, unraveled how LPS is transported to the OM’s surface-exposed outer leaflet by a seven-member (LptABCDEFG) transenvelope complex called “the Lpt system”. (**Fig. 1A**) (18, 19) . Lpt recognizes the lipid A portion of LPS and extracts the molecule from the IM via the LptBCFG transporter, then loading it onto an inter-membrane bridge formed by oligomerized LptA proteins (18, 19). LPS is pushed along the bridge toward the OM LptDE translocon complex that forms a gated pore in the OM which receives LPS and guides its translocation and insertion into the OM outer leaflet (18, 19).

In all *E. coli* K-12 strains, the Lpt system transports only rough LPS since these lab-adapted strains have mutations that inactivated O-Ag biosynthesis (20). Yet, clinical and environmental *E. coli* isolates all produce smooth LPS. As a result, the Lpt system in such bacteria must transport smooth LPS modified with very long O-Ag polysaccharides that can exceed 100 repeat units and 150 nm in length, a formidable challenge in the ∼25 nm width of the periplasm (5, 21). The absence of O-Ag in model *E. coli* K-12 strains means that the impact of smooth LPS production on OM assembly and maintenance of the bacterial cell envelope remains to be fully explored.

In this study, we restored O-Ag biosynthesis in *E. coli* K-12. Surprisingly, the production of O-Ag-modified, smooth LPS presented a significant challenge to commonly used model *E. coli* K-12 strains (such as MC4100 and MG1655), causing broad sensitivity to a several large scaffold antibiotics that are typically excluded by an intact OM barrier (22, 23) . Remarkably, we show that the increased length of the O-Ag polysaccharide compromises OM integrity, causing increased susceptibility to large scaffold antibiotics. Our findings reveal that O-Ag production forces cells into a trade-off: while long O-Ag polymers are critical in the context of infection, their production must be balanced with short O-Ag polymers to avoid significantly compromising the OM antibiotic barrier.

## RESULTS

### Smooth LPS compromises the OM antibiotic barrier of the model *E. coli* K-12 strain MC4100

Integrity of the OM barrier can be assessed by its ability to exclude large scaffold antibiotics (22, 23). Defects in OM assembly or composition allow such antibiotics to permeate into the cell and kill. To probe the effects of smooth LPS production on the OM barrier of the model *E. coli* K-12 strain MC4100, we repaired the *IS5* insertion mutation of its *wbbL* gene, restoring the wildtype allele (*wbbL*^+^) and re-enabling O-Ag synthesis. To our surprise, smooth LPS production caused MC4100 to be sensitive to a range of chemically and functionally distinct large scaffold antibiotics (**Fig. 1B and 1C**). Such antibiotics included the large hydrophilic drug Vancomycin (Van) that targets cell wall synthesis, and the large hydrophobic drugs Erythromycin (Ery) and Novobiocin (Nov) that target protein translation and DNA gyrase/LPS transport, respectively. The broad sensitization to a range of antibiotics with targets throughout the cell was consistent with increased permeability of the MC4100 *wbbL*^+^ OM. The model strain MG1655 (from the same K-12 lineage as MC4100) was also sensitized to antibiotics when producing smooth LPS **(Fig. S1)**.

### Smooth LPS is directly responsible for compromising the OM barrier

In addition to O-Ag, several other cell envelope polymers—peptidoglycan enterobacterial common antigen, and colanic acid—each utilize the Und-P lipid carrier for translocation across the IM (3). We hypothesized that increased OM antibiotic permeability of MC4100 *wbbL*^+^ might be caused by restored O-Ag production limiting Und-P availability for other polymers, thereby broadly impacting the cell envelope and OM. Indeed, prior work found that the Und-P pool becomes limiting if O-Ag is synthesized but smooth LPS production ablated, for example, by deletion of *waaL* (O-Ag ligase) (24). Without WaaL to ligate O-Ag onto LPS, synthesized O-Ag units permanently occupy Und-P, sequestering the lipid carrier and preventing its recycling. Such a scenario leads to severe morphological defects which can be suppressed by simply producing more Und-P via overproduction of UppS (24).

Our MC4100 *wbbL*^+^ cells have a complete O-Ag biosynthetic pathway and make smooth LPS via WaaL. We did not detect any morphological changes caused by *wbbL*^+^ (**Fig. 2A**) and UppS overproduction did not improve the MC4100 *wbbL*^+^ OM barrier against antibiotics (**Fig. 2B**). Hence, our data demonstrated OM antibiotic permeability in MC4100 *wbbL*^+^ is not caused by Und-P limitation. Underscoring that conclusion, we found that Δ*waaL* in fact strongly suppressed OM antibiotic permeability of MC4100 *wbbL*^+^ (**Fig. 2C**). This suppression is remarkable since it occurred despite Δ*waaL* also causing the expected morphology (cell length and width) defects in *wbbL*^+^ that prior studies attributed to Und-P limitation (**Fig. 2A**). Our Δ*waaL* suppressor data pinpoint the culprits of OM permeability in MC4100 *wbbL*^+^: it is the smooth LPS molecules themselves. Lacking WaaL, smooth LPS cannot be produced. Surprisingly, an OM composed of rough LPS forms a better antibiotic barrier than an OM composed of smooth LPS.

**Fig. 2.**
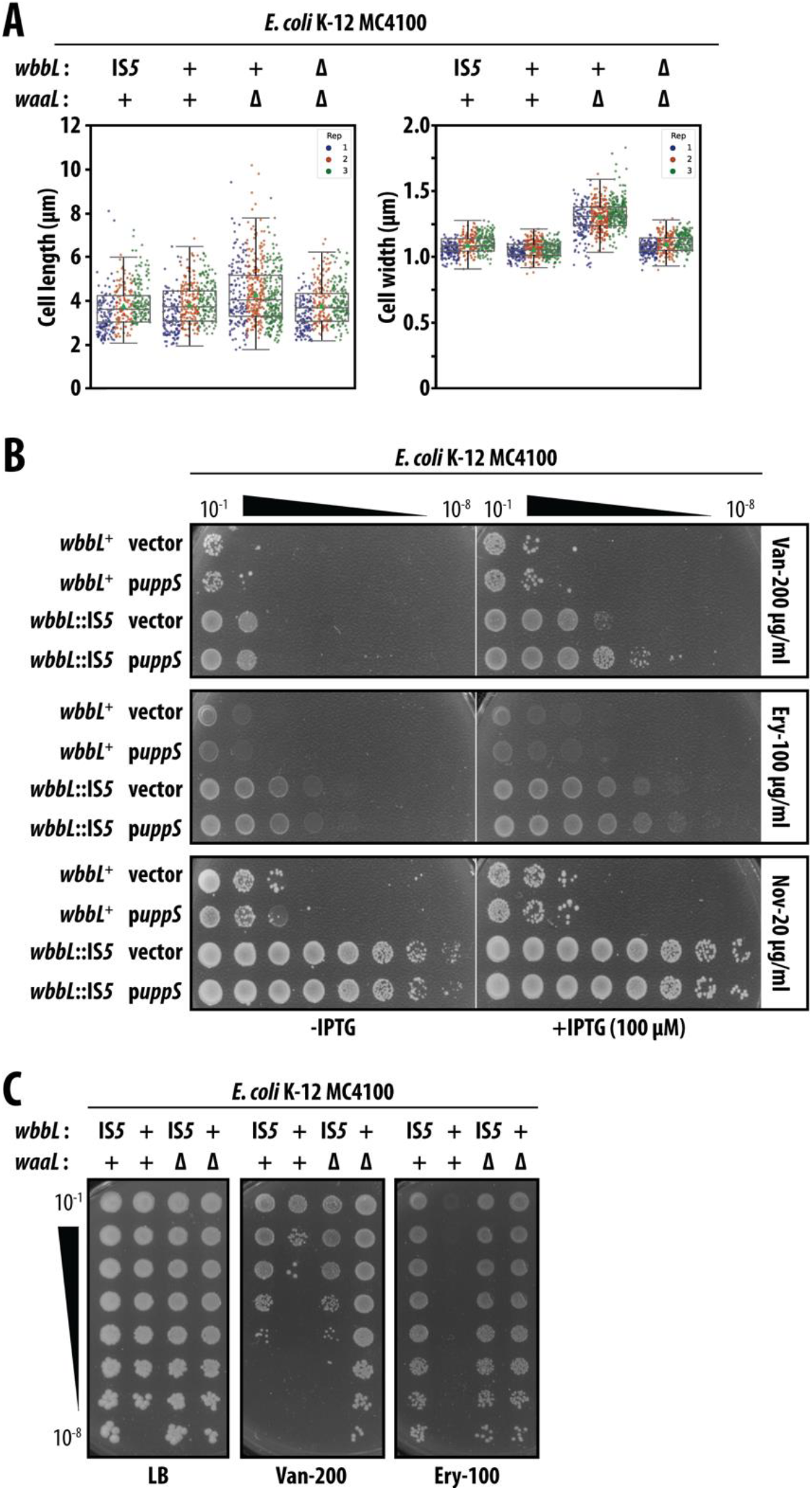
OM antibiotic permeability of smooth LPS producing MC4100 is not due to sequestration of the Und-P lipid carrier. **(A)** Measurement of cell length and cell width morphology parameters. Data from three replicate assays presented, mean ± standard deviation shown. **(B)** Efficiency-of-plating antibiotic sensitivity assays using 10-fold serial dilution of saturated cultures. IPTG is an inducer of plasmid-encoded UppS. **(C)** Antibiotic sensitivity assays using 10-fold serial dilution of saturated cultures. All data are representative of at least three independent experiments.

### Analysis of LPS transport and OM properties of smooth LPS strain of MC4100

We reasoned that smooth LPS molecules may be less efficiently transported to the OM by the Lpt pathway than rough LPS molecules, given their added long, complex O-Ag polysaccharide. To examine LPS content in cellular membranes, we lysed cells, separated IM and OM fractions in sucrose density gradients, and compared the abundance of LPS in each fraction. Examining the distribution of all LPS forms between IM and OM fractions, we did not detect major differences but did note that MC4100 *wbbL*^+^ had a minor increase in IM LPS content and a concomitant minor decrease in LPS within OM fractions compared to its MC4100 *wbbL*::IS*5* parent (**Fig. S2A**).

In *E. coli*, it is well established that decreased LPS content in the OM is offset by translocation of OM inner leaflet PLs to the outer leaflet, leading to a loss of lipid asymmetry in the OM bilayer (25–27). Such PL mislocalization sensitizes cells to detergents such as sodium dodecyl sulfate (SDS) and bile (25, 28). Loss of lipid asymmetry can be exacerbated by chelating agents (e.g. EDTA) which disrupt LPS packing and promote further PL mislocalization (25, 27, 28). We used such phenotypic assays to assess *wbbL*^+^ relative to well-characterized mutations impacting the OM lipid bilayer (**Fig. S2B**). As expected, mutations that disrupt LPS transport (*lptFG-*depletion) exhibited sensitivity to both bile and SDS-EDTA. Similarly, inactivation of the Mla-PldA lipid asymmetry maintenance system (Δ*mlaC* Δ*pldA*), which has the effect of allowing mislocalized PLs to accumulate in the outer leaflet, also caused the expected bile and SDS-EDTA sensitivity (25, 28). Smooth LPS producing MC4100 *wbbL*^+^ cells were not appreciably sensitized to either bile or SDS-EDTA, compared to the rough LPS producing MC4100 (*wbbL*::*IS5*) parent (**Fig. S2B**). Remarkably, Mla system inactivation (Δ*mlaC*) in MC4100 *wbbL*^+^ had no appreciable impact on further sensitizing these smooth LPS producing cells to large scaffold antibiotics (**Fig. S3A**).

Our data demonstrated that MC4100 *wbbL*^+^ cells phenotypically exhibited no signs of significant broken OM lipid asymmetry. Hence, the minor difference in LPS distribution between IM and OM that we detected in MC4100 *wbbL*^+^ appeared unlikely to explain the OM barrier defect that causes antibiotic sensitivity in *wbbL*^+^.

A previous study reported that smooth LPS production increases the stiffness of the OM (29). We considered that increased OM stiffness may impact antibiotic barrier function in MC4100 *wbbL*^+^. Hence, we tested whether *wbbL*^+^ antibiotic sensitivity could be alleviated by deleting *ompA* (encoding a major porin that links OM and PGN cell wall) as this mutation was reported to significantly decrease OM stiffness in that same study (29). However, Δ*ompA* failed to suppress OM antibiotic permeability of MC4100 *wbbL*^+^ (**Fig. S3B**), indicating that the *wbbL*^+^ OM barrier does not benefit from reducing stiffness and that a stiff OM is unlikely to be the cause of antibiotic permeability of MC4100 *wbbL*^+^.

### Shortening O-Ag polysaccharides suppresses antibiotic sensitivity

To better understand the basis of the compromised OM antibiotic permeability barrier in MC4100 *wbbL*^+^, we designed a genetic selection strategy to recover spontaneous mutations that suppress vancomycin sensitivity when smooth LPS is produced. In our selection strategy, we employed rough-specific Autographiviridae Ffm bacteriophage to select against mutations that simply inactivated O-Ag synthesis or prevented O-Ag attachment to LPS, as we already knew such mutations suppressed vancomycin sensitivity (e.g. nulls of *wbbL* or *waaL*).

One informative suppressor caused a Q205Am nonsense mutation in the 326 amino acid WzzB protein, the O-Ag chain length regulator (**Fig. 3A**). This mutation was a null allele since we found that introducing a Keio Δ*wzzB*::*kan* deletion-insertion allele into MC4100 *wbbL*^+^ also suppressed vancomycin sensitivity of MC4100 *wbbL*^+^ (**Fig. 3B**) Inactivation of *wzzB* effectively restored the OM barrier against not only vancomycin, but also erythromycin and novobiocin (**Fig. 3A**). Hence, the *wzzB* mutation broadly improved the OM antibiotic barrier.

**Fig. 3.**
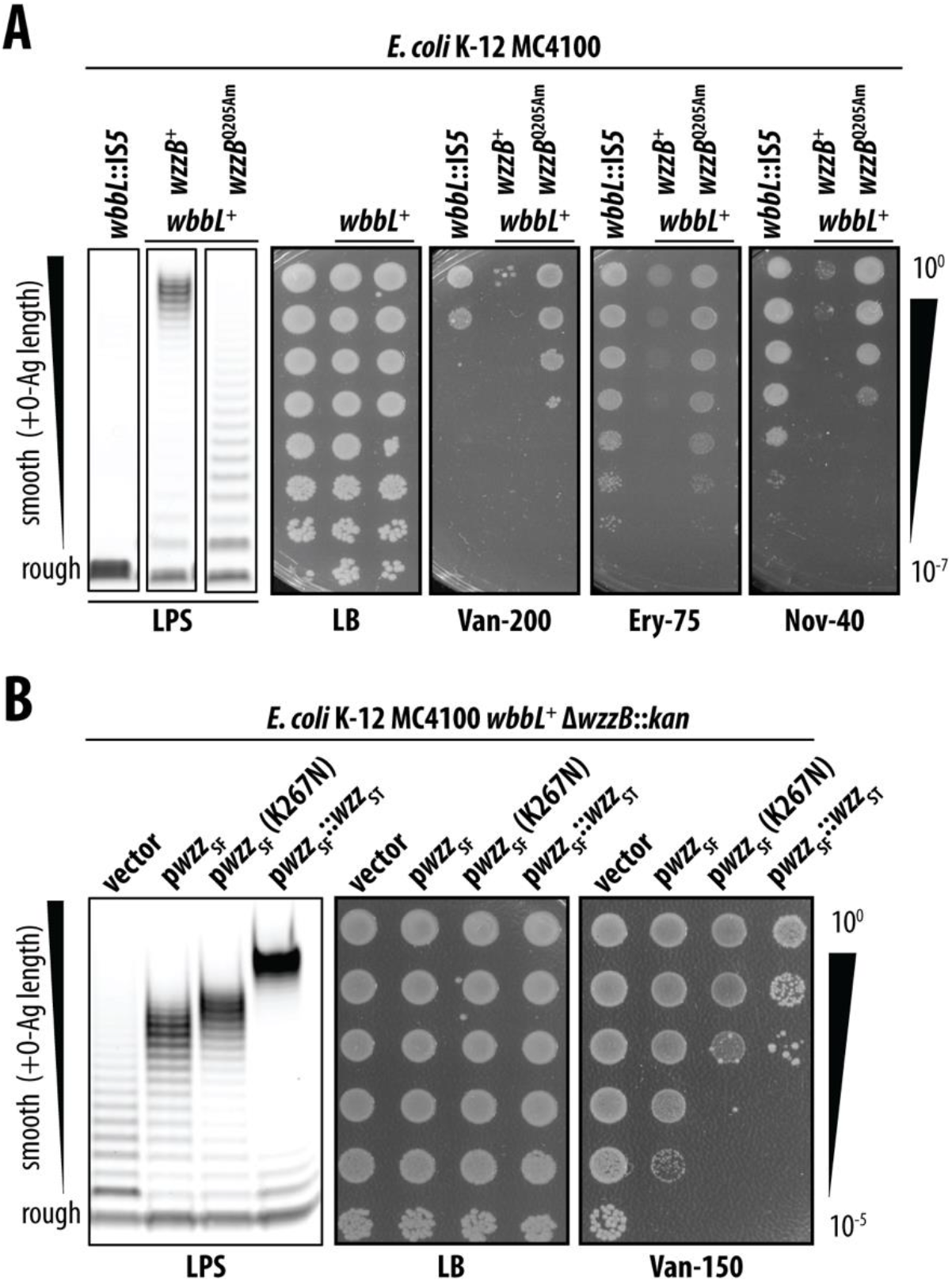
O-Ag polysaccharide length impacts OM antibiotic permeability. **(A)** Whole-cell LPS profiles for *E. coli* strains and corresponding antibiotic sensitivities assessed by efficiency-of-plating assays. *wzzB*^Q205Am^ is a suppressor of *wbbL*^+^ antibiotic sensitivity. **(B)** Whole-cell LPS profiles and corresponding vancomycin sensitivity for *E. coli* strains producing varying *wzzB* alleles *in trans*. Ten-fold serial dilutions of saturated cultures are shown. Antibiotic concentrations are in μg/ml. All data are representative of at least three independent experiments.

MC4100 *wbbL*^+^ cells with wildtype *wzzB*^+^ produce smooth LPS molecules that have been modified with O-Ag polysaccharides of varying lengths, but which have a characteristic modal length of ∼16 O-Ag repeat units. *wzzB* inactivation led to an unregulated O-Ag chain length and a characteristic continuous ladder of smooth of LPS lengths that gradual decreases in abundance at each successive chain length. Overall, the effect of the *wzzB* suppressor was for cells to produce smooth LPS molecules that are much shorter than the long smooth LPS produced by *wzzB*^+^ cells. Apparently, shortening the smooth LPS molecules can restore the OM antibiotic barrier.

### Lengthening O-Ag polysaccharides increases antibiotic susceptibility

Our data suggested that, in *E. coli* K-12 MC4100, smooth LPS impairs the OM permeability barrier and that producing shorter smooth LPS can improve barrier integrity.

We next sought to directly test whether altering the length of smooth LPS molecules would correspondingly alter OM permeability in *E. coli* MC4100 *wbbL*^+^. To manipulate smooth LPS length, we complemented a Δ*wzzB* mutation in MC4100 *wbbL*^+^ with a plasmid-encoded *wzzB*_SF_ from *S. flexneri* (**Fig 3B**). The resulting strain produced short-type smooth LPS (10-16 repeats) and was only very modestly sensitized to vancomycin (**Fig 3B**). Providing a plasmid encoding a K267N variant of Wzz_SF_ (that increased O-Ag chain modal length to ∼14-18 repeat units) had the expected effect of lengthening the smooth LPS and this exacerbated vancomycin sensitivity (**Fig 3B**). Providing a plasmid encoding a synthetic WzzB hybrid of *S. flexneri* and *Salmonella* serovar typhi orthologs further lengthened the smooth LPS produced (to a model length of >20 O-Ag repeat units) and further still increased vancomycin sensitivity (**Fig 3B**). These data directly link longer LPS O-Ag production with an impaired OM antibiotic permeability barrier.

### Increased rough or short smooth LPS production improves OM barrier integrity

The widely studied model strains of *E. coli* K-12 MC4100 and MG1655 both belong to the “EMG2” lineage of *E. coli* K-12 and share the *wbbL*::*IS5* mutation that inactivated their O-Ag biosynthetic pathways. Another, independent, lineage of *E. coli* K-12 also exists, named “WG1”, to which belongs the strain NCM3722. Among other genetic differences, WG1 strains lost production of O-Ag due to a different mutation. In WG1 strains, *wbbL* is intact but a large deletion of the *rfb* O-Ag biosynthetic locus causes rough LPS production. We repaired the *rfb* locus to yield NCM3722 *rfb*^+^ in which O-Ag production is restored and smooth LPS is produced. Strikingly, in contrast to strains of the EMG2 lineage, NCM3722 *rfb*^+^ exhibited much less smooth LPS dependent antibiotic sensitivity (**Fig. 4A)**. Similarly, in *Shigella flexneri* 2457T, a human pathogen that is closely related to *E. coli* K-12 strains (and which naturally retained O-Ag synthesis), smooth LPS production did not cause a dramatic increase in antibiotic sensitivity compared to a rough LPS producing Δ*rmlD*::*kan* mutant (inactivated for O-antigen biosynthesis) (**Fig. 4A**). Importantly though, in both NCM3722 and *S. flexneri*, the production of rough LPS was nonetheless better at resisting large scaffold antibiotics (**Fig. 4A**). Apparently, even for a pathogen such as *S. flexneri*, production of rough LPS improves integrity of the OM antibiotic permeability barrier.

**Fig. 4.**
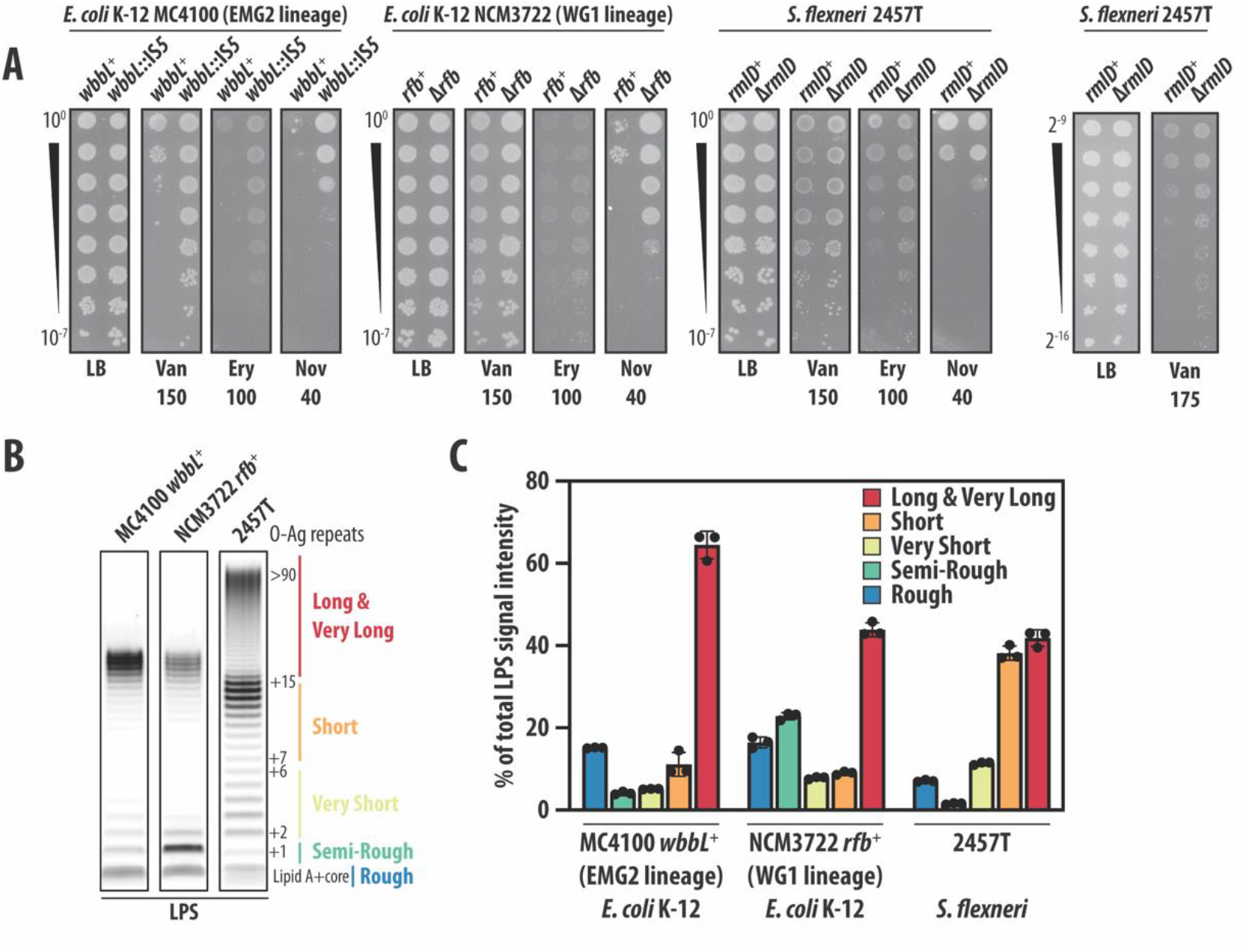
An increased proportion of rough, semi-rough, or short smooth LPS increases OM antibiotic barrier integrity. **(A)** Efficiency-of-plating assays assessing antibiotic sensitivity of rough and smooth LPS producing strains. Rough LPS producers are *wbbL*::IS*5*, Δ*rfb*, Δ*rmlD*, in MC4100, NCM3722, and 2457T, respectively. Ten-fold serial dilutions of saturated cultures shown. Antibiotic concentrations are in μg/ml. **(B)** LPS profiles of smooth *E. coli* (MC4100 and NCM3722) *and Shigella* strains. (**C**) Quantification of LPS profile distribution in each strain. Data are means ± standard deviation of LPS band intensities from triplicate Emeral ProQ stained LPS gels, relative to total LPS. All data are representative of at least three independent experiments.

We noted that both NCM3722 and *S. flexneri* encode additional, paralogous copies of LptF and LptG LPS transport proteins whereas MC4100 (and the EMG2 lineage) do not. However, this genetic difference did not account for the difference in OM antibiotic permeability when smooth LPS is produced (**Fig. S4)**.

Analysis of the distribution of smooth LPS lengths in *E. coli* MC4100 and NCM3722 and *S. flexneri* 2457T demonstrated that MC4100 produces mostly smooth LPS molecules >16 O-Ag repeat units which is considered a long-type LPS (30) (**Fig. 4B**). In contrast, while O-Ag chain length in *S. flexneri* 2457T is bimodal, consisting of both very long (>90 O-Ag repeat units long) and short LPS (<16 O-Ag repeat units long), *S. flexneri* primarily produces the short-type smooth LPS molecules (**Fig. 4B and 4C**) determined by a WzzB ortholog (Wzz_SF_) (31). Moreover, although NCM3722 and MC4100 encode the same WzzB, NCM3722 produces strikingly more LPS that has been modified by unpolymerized O-Ag (only a single O-Ag repeat unit). Therefore, the majority of NCM3722 smooth LPS molecules are also shorter, compared to MC4100 (**Fig. 4B and 4C**). Collectively, our data leads us to a model where smooth LPS molecules can impair the OM antibiotic barrier, and that longer smooth LPS molecules impair the OM more than short smooth LPS molecules. Hence, OM integrity in smooth LPS producing bacteria relies on cells balancing production of short and long smooth LPS. Indeed, our data suggested that even a modest shift in the balance of long and short or rough LPS O-Ag can greatly impact OM permeability.

To test our model in MC4100 *wbbL*^+^, we used heterologous expression of the *waaL* O-Ag ligase from *S. flexneri* (*waaL*^SF^). WaaL^SF^ is specific for the *S. flexneri* “R3” core oligosaccharide and does not recognize the “K-12” core oligosaccharide of MC4100, hence WaaLSF cannot attach O-Ag to MC4100 LPS; however, WaaL^SF^ can recognize the undecaprenol pyrophosphate (Und-PP)-linked O-Ag polymers produced by in MC4100 *wbbL*^+^ (3, 32–34). Hence, we reasoned that heterologous production of WaaL^SF^ would interfere with smooth LPS production by limiting the amount of O-Ag that is available to the native MC4100 WaaL for ligation onto LPS. Indeed, expression of this foreign O-Ag ligase in MC4100 *wbbL*^+^ had the effect of increasing the proportion of rough LPS at the cost of the long smooth LPS produced (**Fig. 5**). This change in LPS profile was sufficient to suppress MC4100 *wbbL*^+^ sensitivity to vancomycin and other large scaffold antibiotics (**Fig. 5**). This finding underscored our model that maintaining a balanced proportion of rough (and short) LPS ensures OM antibiotic barrier integrity when producing smooth LPS.

**Fig. 5.**
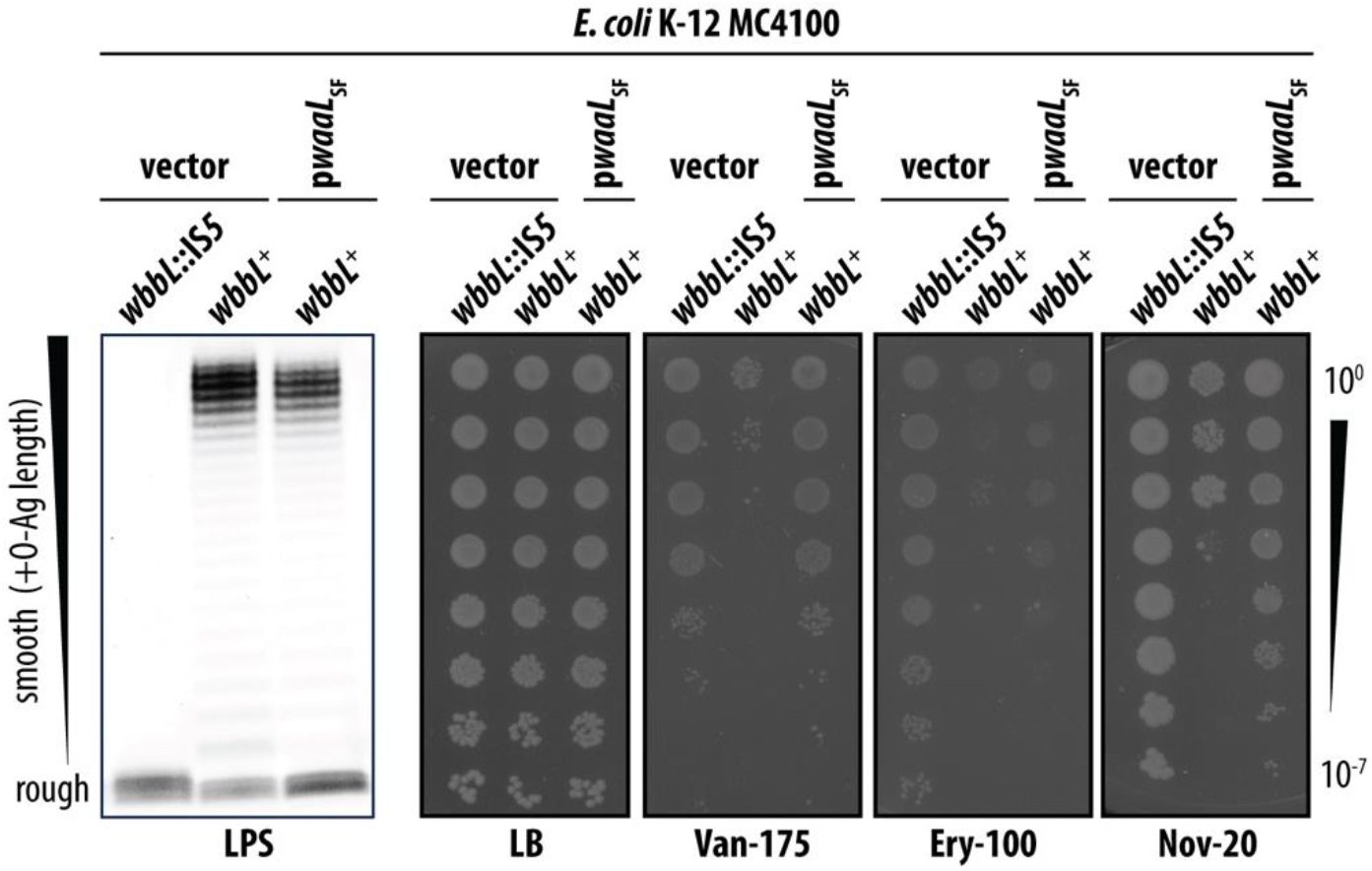
Expression of *S. flexneri* WaaL suppresses antibiotic sensitivity of O-Ag producing *E. coli* MC4100 by altering the proportion of rough and smooth LPS molecules produced. LPS profiles of MC4100 expressing non-native *S. flexneri* O-Ag ligase WaaL_SF_ (p*waaL*_SF_) showing an increased abundance of rough LPS population at the expense of a reduction in smooth LPS population. Increased proportion of rough LPS improves the OM antibiotic barrier of MC4100 *wbbL*^+^ expressing WaaL_SF_. All data are representative of at least three independent experiments.

## DISCUSSION

There is a pressing need for a detailed understanding of how molecules penetrate across the OM so that this information can be leveraged for designing novel antibiotics. Most studies of OM biogenesis and LPS transport have exploited the genetic tractability of *E. coli* K-12 lab-adapted strains which inherently produce only rough LPS. Remarkably, we discovered that the *E. coli* OM is permeabilized for antibiotic entry by the production of O-Ag polysaccharides that decorate smooth LPS molecules, which represent the natural LPS forms in clinical and environmental *E. coli*. Indeed, we found the human pathogen *S. flexneri* also has a more permeable OM when producing smooth LPS and that its OM antibiotic barrier can be enhanced when producing only rough LPS (following inactivation of O-Ag synthesis). Surprisingly, our data lead to us to a model where long smooth LPS molecules compromise the OM barrier and rough (or short) LPS improve the OM barrier.

Studies of enteric bacteria across decades have cataloged ways in which regulation of O-Ag polysaccharide length is critical for bacteria to survive within a mammalian host and to enact their virulence program (5). The length of long O-Ag polysaccharides must be controlled so that bacteria are shielded from host assaults, but cell surface virulence factors can still readily access host targets. Our findings reveal that an unappreciated benefit for this regulation is the cell’s need to maintain the intrinsic impermeability of its OM. A pathogen such as *S. flexneri* appears to balance these competing needs by producing generally short smooth LPS (<17 O-Ag repeats) and this allows its OM to be remarkably robust against large-scaffold antibiotics. Similarly, the WG1 lineage *E. coli* K-12 strain NCM3722 produces an abundance of LPS capped with only a single O-Ag unit (termed “semi-rough” LPS) such that the OM permeabilizing effect of producing very long or long smooth LPS is offset by simultaneously producing a large population of rough, semi-rough, and short LPS. In contrast, strains of the major EMG2 *E. coli* K-12 lineage (e.g. MC4100 and MG1655) commit the majority of LPS to the long smooth form, and this makes them susceptible to antibiotics.

In the context of bacterial pathogenesis, virulence and phage resistance, the vulnerability of mutants producing only rough LPS has also long-been acknowledged(5, 17, 35); such molecules are not helpful in this context. Yet, curiously, it was also long evident that *E. coli, S. flexneri*, and other Enterobacterales produce large populations of rough (and short) LPS forms - why this occurs has been unclear. We propose that a benefit of co-producing rough (and short) LPS O-Ag forms is so that bacteria can maintain the integrity of their OM barrier.

To be clear, we do not propose that smooth LPS producing bacteria— commensals or pathogens—inherently produce a permeabilized OM that causes them to be susceptible to large scaffold antibiotics. Enterobacterales are not clinically vulnerable to vancomycin therapy. We find that the OM barrier of *S. flexneri* can be improved by rough LPS production, but only modestly. Hence, we suggest that native smooth LPS producers (such as *S. flexneri*) have evolved to effectively balance the production of different smooth LPS molecules to efficiently optimize for two parameters: host pathogenesis and OM barrier integrity.

Other Gram-negatives, including strains of *E. coli* and *Klebsiella pneumoniae*, lack a periplasmic WzzB O-Ag length regulator and instead use an entirely distinct, cytosolic mechanism to establish a modal O-Ag length (6). In such bacteria, the O-Ag polysaccharide is first assembled at the cytoplasmic face of the inner membrane and then flipped across the IM by an ABC transporter(6). Despite differing modes of O-Ag synthesis, it is notable that such bacteria also exhibit a balanced distribution of smooth LPS O-antigen lengths. For example, *E. coli* O9a and *K. pneumoniae* O12 exhibit a tight unimodality with nearly half the O-Ag polysaccharides being 13-14 repeat units or less (6). Meanwhile, *K. pneumoniae* O2a and O1 exhibit a wider distribution of O-Ag lengths that include longer polysaccharides, but, markedly, this is accompanied by a considerable amount of rough LPS (6). An interesting implication of our model is that there may be an inherent benefit to producing LOS, lacking any O-Ag polysaccharide, for tight OM barrier integrity and hence resistance to antibiotics or environmental toxins. We speculate that this previously unappreciated advantage of LOS may be the reason why production of LOS is highly prevalent among diverse Gram-negative species.

The fact that smooth LPS has a permeabilizing effect on the OM is underscored by D*waaL* suppression of *E. coli* MC4100 *wbbL*^+^ antibiotic sensitivity. D*waaL* potently increased resistance to functionally distinct large scaffold antibiotics simply by preventing smooth LPS from being formed and, remarkably, this suppression was effective despite D*waaL* clearly also causing serious defects in the cell envelope (reflected by morphological abnormality). Recently, Qin et al (36) reported that restoring O-Ag to K-12 sensitized cells to the combination treatment of bile with vancomycin. Since weakening the cell-wall is known to sensitize cells to both detergents such as bile and cell wall-targeting antibiotics such as vancomycin, the authors suggested that O-Ag synthesis results in a deprivation of sufficient Und-P for efficient cell wall synthesis. Indeed, the authors discounted a possible effect of O-Ag production on OM permeability. Our data challenges their proposal in several ways. Firstly, we measured no obvious morphological changes in MC4100 *wbbL*^+^ cells that would be expected upon Und-P sequestration. Secondly, UppS overexpression (increasing Und-P availability) failed to suppress smooth LPS producing MC4100 *wbbL*^+^ sensitivity to vancomycin (or other large scaffold antibiotics). Moreover, we find that smooth LPS production sensitizes *E. coli* (and *S. flexneri*) to not only vancomycin but to a range of large scaffold antibiotics that target distinct cellular pathways that do not rely on Und-P; these antibiotics all share the property of being effectively excluded from the cell by an intact OM barrier. Finally, we found that the sensitivity of smooth LPS producing *E. coli* to vancomycin and the other antibiotics can be potently suppressed by deleting *waaL*, preventing the ligation of O-Ag polysaccharide to the LPS core. Suppression by D*waaL* is effective even though it is known to actively cause severe Und-P sequestration (since, absent WaaL, the Und-P-attached O-Ag intermediate has no way to be liberated from Und-P and ligated onto LPS) (24). Collectively, our data provide several lines of evidence that the vancomycin sensitivity caused by smooth LPS does not stem from Und-P limitation.

We detected a minor defect in LPS distribution between IM and OM, suggesting that smooth LPS production may result in a minor impairment in Lpt transport efficiency. Previous *in vitro* experiments showed that the lipid A portion is sufficient for LptA binding and that O-Ag does not impact this interaction (37). Hence, it is likely that the LptA bridge accommodates either smooth or rough LPS without distinction. *In vitro* reconstitution studies suggested that changes in the core oligosaccharide affect transport efficiency at LptBFG (38); this could be due either to altered substrate recognition by LptBFG or altered physical properties of LPS that increase its tendency to aggregate and limit diffusion in the transporter (38, 39). It is possible that the same lateral interactions between adjacent LPS O-Ag polysaccharides that have been proposed to increase stiffness and strength at the OM also increase the tendency for LPS to aggregate in the IM and influence transport efficiency. In any case, any Lpt transport deficiency in smooth LPS-producing cells is likely minimal as it clearly does cause hallmarks of broken OM lipid asymmetry (such as detergent sensitivity) which occur if LPS were significantly limiting in the OM. In all, we do not think that any impacts on transport via Lpt explain the disrupted OM antibiotic barrier integrity caused by smooth LPS.

It is notable that passage of large antibiotics is thought to occur through β-barrels of some export systems. In the most well explored example, vancomycin is thought to pass through the lumen of the TolC OM β-barrel which, with its partners AcrAB, forms the Type I secretion system for antibiotic efflux. Indeed, *E. coli* mutants lacking TolC are up to 4-fold more resistant to vancomycin. Since TolC-dependent vancomycin-sensitivity also requires AcrAB, it seems that vancomycin can only pass through the OM via TolC when the protein is actively engaged by the AcrAB pump (40, 41). It has been suggested that other OM systems, like the Bam complex, offer additional avenues for antibiotics to enter the cell (42). By analogy, it may be that the LptD-LptE complex may offer another entryway through the OM into the cell for large antibiotics. LptD is a very large 26-strand β-barrel protein with a sizeable hydrophilic cavity. Extracellular loops of LptD fold into the barrel lumen, sealing off the cavity. The LptE partner lipoprotein extends into the LptD lumen to act as a plug (43–46). An LptE variant defective in plugging sensitizes cells to large-scaffold antibiotics, demonstrating that a poorly plugged LptD-LptE provides a ready conduit for antibiotic entry into the cell (47). Conformational changes in LptD-LptE that open the barrel are likely to occur during LPS passage through the translocon. It is tempting to speculate that the impact of O-Ag on OM permeability is that smooth LPS substrates alter the translocon in a way that enables antibiotics to pass more readily through the LptD barrel, perhaps by keeping the LptD-LptE complex in an opened conformation for an extended period while the long O-Ag polysaccharide is extruded. Indeed, longer lengths of O-Ag polysaccharide attached to LPS lead to increased antibiotic sensitivity.

Ultimately, our findings imply that enteric bacteria are faced with a balancing act when producing LPS: long smooth LPS will protect against the host, but permeabilize the OM; rough or short smooth LPS endangers survival in a host but ensures a robust OM barrier. Cells need to ensure balanced production of the different LPS types in order to thrive.

## MATERIALS AND METHODS

### Bacterial strains, plasmids, and growth conditions

Strains, plasmids, and oligos used in this study are listed in Tables S1-S3. Isogenic derivative strains were constructed by P1vir transduction or λRed recombineering (in *S. flexneri* 2457T and *E. coli* NCM3722). To generate MC4100 cells that produce smooth LPS, the *wbbL*::IS*5* mutation was repaired by transducing the *wbbL*^+^ locus from NCM3722. To generate NCM3722 cells, the large deletion of its *rfb* locus was repaired by transducing *rfb*^+^ from MC4100. In all cases, transductants producing smooth LPS were selected by their resistance against Ffm phage, a rough LPS-specific bacteriophage (48). The resulting smooth LPS producing strains were confirmed by whole-genome sequencing (WGS); they were confirmed to have the expected wildtype sequences at *wbbL* or *rfb* (in MC4100 and NCM3722, respectively) and lack additional mutations in comparison to their corresponding rough LPS producing parental strains. WGS data is deposited in the NCBI Sequence Read Archive (SRA) under BioProject PRJNA1304456. Keio deletion-insertion alleles or Tn*10* transposon insertions were used (49, 50). Introduction of Keio alleles and the presence of suppressor mutations were routinely verified by PCR and Sanger sequencing.

Strains were cultured in LB (Lennox) broth or agar at 37°C. Media were supplemented with ampicillin (100 µg/mL), chloramphenicol (20 µg/mL), kanamycin (25 µg/mL), tetracycline (25 µg/mL), as required for marker selection.

### Antibiotic sensitivity assays

Efficiency of plating assays were used to determine the relative sensitivities of strains to various treatments. Serial ten-fold dilutions of saturated cultures (standardized by OD_600_) were replica-plated onto LB agar media and incubated overnight at 37 °C. Time-kill assays diluted log-phase cultures to OD_600_ of 0.1 into 0.2ml media per well of a 96 well plate. Cells were grown at 37°C with linear shake at 567 cpm (3 mm) and OD_600_ was measured every 10 min using a BioTek Synergy H1.

### Suppressor selection

Spontaneous suppressor mutations of MC4100 *wbbL*^+^ vancomycin sensitivity were isolated on LB media supplemented with both vancomycin and rough-specific Ffm phage. The *wzzB*(Q205Am) suppressor mutation was identified by WGS and independently confirmed by Sanger sequencing. WGS data is deposited in NCBI SRA under BioProject PRJNA1304456.

### Microscopy and image analysis

Cells in mid-exponential growth (OD_600_ ∼0.2) were placed between a no.1.5 cover glass and 1mm-thick 1% agarose pad made with LB Lennox. Cells were imaged using an inverted fluorescence microscope (Olympus IX83) with an oil immersion phase-contrast 60x objective. Images were acquired using a Neo 5.5 sCMOS camera (Andor) and MetaMorph software (Molecular Devices). Cell shape measurements were performed using MicrobeJ 5.13o(4)(53), a plug-in for the ImageJ/Fiji software(54).

### Sucrose density gradient fractionation

Separation of membranes by sucrose density gradient fractionation was performed as described previously (55). Briefly, sub-cultures of each strain were grown in LB broth with aeration until O.D. of 0.6–0.8. Cells were washed with cold 10mM Tris-HCl, pH 8.0, and centrifuged at 10 000 x g for 10 min at 4°C. Cells were resuspended in 20 ml of 10 mM Tris-HCl, pH 8.0 containing 20% sucrose (w/w), Benzonase (EMD Millipore) and HALT protease inhibitor cocktail (Thermo Scientific), and then lysed with a single passage through a French Pressure Cell Press (Thermo Spectronic) at 8000 psi. Unbroken cells were removed by centrifugation at 10 000 x *g* for 10 min at 4°C. The cleared cell lysate was collected, and 5.5 ml was layered on top of a two-step sucrose gradient consisting of 5 ml of 40% (w/w) sucrose solution layered on top of 1.5 ml of a 65% (w/w) sucrose solution in an Ultra-Clear tube (14 x 89mm, Beckman Coulter). All sucrose solutions were prepared in 10mM Tris-HCl, pH 8.0. To separate the inner and outer membranes, samples were centrifuged at 35 000 rpm for 18 h in a Beckman SW41 rotor in an OptimaXE-90 Ultra-centrifuge (Beckman Coulter). Twelve fractions (of 1.1 ml) were manually collected from each tube, starting from the lowest density i.e. from the top of the tube.

### LPS profiles and analyses

LDS sample buffer (Life Sciences) with 4% β-mercaptoethanol was added to either sucrose gradient fractions or whole cell pellets consisting of 5 × 10^8^ cells from an overnight culture. Samples were denatured at 100°C for 10 min, allowed to cool, and then treated with 125 ng/μl proteinase K (New England Biolabs) at 55°C for 16 h. Proteinase K was heat inactivated, and the lysates were resolved by SDS-PAGE. Gels were stained with the Pro-Q Emerald 300 LPS Gel Stain kit (Molecular Probes) in accordance with the manufacturer’s instructions. LPS bands were visualized by UV transillumination. Each band within a lane represents a population of LPS with a discrete O-Ag polysaccharide length. The relative band intensities were determined with the Quantity One imaging software (Bio-Rad). Combined intensities defined the total LPS of samples. Individual bands or band groupings of rough, semi-rough, very short, short, and long with very long LPS types were measured and quantified as a percentage of total LPS.

### Western immunoblotting

Sucrose gradient fractions were diluted in 2x LDS Sample Buffer (Life Sciences) with 4% beta-mercaptoethanol (β-ME) and incubated at 100°C for 5 min. Samples were resolved by polyacrylamide gel electrophoresis on 4-12% Bis-Tris gels (Novex). Resolved proteins were transferred to 0.2 μm nitrocellulose membranes and probed with anti-LptD polyclonal antisera (Silhavy Lab stock), used at a dilution of 1:5,000. The anti-IMP/LptD polyclonal antibody also reacts with a 55-kDa inner membrane protein and the OmpA (37 kDa) outer membrane protein (56). Membranes were subsequently probed with goat anti-rabbit-HRP secondary antibodies (EMD Millipore). Probed membranes were developed by incubating with Immobilon Classico Western HRP substrate (EMD Millipore). The resulting chemiluminescence was detected with a BioRad ChemiDoc MP. The relative band intensities were determined with the Quantity One imaging software (Bio-Rad) and the intensity of the band in each fraction expressed as a percentage of the total cumulation of bands across that gradient.

## ACKNOWLEDGMENTS

This work was supported by NIH grants R35 GM156739 (to MG), U19 AI158080 (to MK), and R25 AI175048 (supporting BGP). We are grateful to Kevin Young for plasmids pDSW204 and pDSW204-uppS, and Renato Morona for the suite of plasmids expressing *wzzB* variants. We thank Taryn Trigler for helpful comments on the manuscript.

